# TriScan: High-Contrast Confocal Sectioning with Widefield Speed

**DOI:** 10.1101/2023.04.11.536163

**Authors:** Robin Van den Eynde, Jon Verheyen, Riley Hargreaves, Vincent Van Deuren, Damla Temel, Adrián Gómez Sánchez, Timon Vlems, Paul Miclea, Qendresa Arifi, Hélène Roberge, Frederik Niekamp, David Boulet, Marta Ciwinska, Colinda Scheele, Olivier Devos, Alessandro Furlan, Thomas Huser, Mark Bates, Marcel Leutenegger, Theo Lasser, Sebastian Munck, Lydia Danglot, Cyril Ruckebusch, Josef Lazar, Wim Vandenberg, Peter Dedecker

## Abstract

Modern analytical methods increasingly require fast, high-resolution, and scalable 3D fluo-rescence imaging with optical sectioning. We present the TriScan, a microscope that allows for compact, inexpensive, and easy-to-implement confocal imaging at very fast imaging speeds. The TriScan combines line-scan confocal imaging with a sensitive camera detector via a simple optical layout that imposes no practical limit on the scanning rate. By combining this method with fast resonant scanners, the TriScan offers the speed and convenience of widefield imaging but adds confocal sectioning, offering a friendly and familiar front-end that is easy to operate. We have developed multiple versions of the TriScan that can be added to existing microscopes, with all scanner, laser and camera timing handled by the modules themselves. We demonstrate the versatility of the system across multicolour, 2D, 3D, dynamic, and super-resolution imaging. By combining high performance with a simple and robust layout, the TriScan can support a broad range of applications ranging from research to diagnostics and screening.

## 1 Introduction

Fluorescence microscopy is an indispensable tool in the life and material sciences, driven by its combination of molecular specificity, very high sensitivity, low invasiveness, and compatibility with a very broad range of samples. Over the past years, the technique has become a key component in a wide range of advanced workflows, including approaches such as in situ transcriptomics [1], multiplexed immunohistochemistry [2], high-content phenotypic screening [3] and organ-on-chip imaging [4]. Performing these in the structurally intact (3D) samples allows preservation of the native spatial organization and relationships between cells and tissues, better approximating the native conditions and preventing the loss of information that can occur when samples are sectioned or otherwise disrupted. The high potential of these methods for applications within areas such as precision medicine supports that advanced fluorescence imaging will become a standard tool in many clinical and diagnostic workflows.

To succeed, such imaging needs to offer not only sophisticated imaging performance, including throughput, sensitivity, and resolution, but also needs to achieve this in a format that is simple to operate, robust, and cost-effective. For imaging of 2D flat samples, this challenge can be solved using widefield (camera-based) imaging, the gold standard for speed, sensitivity, and simplicity. However, achieving this in 3D imaging has been much more challenging because widefield imaging does not provide optical sectioning.

Several methods have been proposed to address this challenge. Confocal imaging introduces an optical pinhole to reject out-of-focus light. However, this method’s throughput is limited by the need for point-by-point scanning, requiring complex control and synchronization to function well. One solution is to parallelize the process using spinning disk confocal microscopes, which illuminate and detect from an array of pinholes simultaneously, though the systems are optically and mechanically complex, suffer from pinhole cross-talk and limited light throughput, and can be difficult to integrate with advanced detection modalities such as flexible spectral imaging [5].

Other approaches modify the optical geometry of the system, including light-sheet fluorescence microscopy. These combine widefield detection with illumination confined to a thin sheet of light, achieving rapid volumetric imaging while minimizing photobleaching and phototoxicity. However, classical implementations require an orthogonal multi-objective geometry that complicates the use of standard slides and multi-well plates, and limits the numerical aperture of the imaging. Various strategies have been proposed to overcome these limitations, including the use of mirrored sample holders [6], oblique light sheet illumination and detection [7–10], and custom optical elements. These innovations enable high-performance imaging but increase system complexity, cost, and alignment requirements, reducing suitability for routine applications such as screening and diagnostic imaging. Furthermore, their optical layouts are intrinsically geared towards deep optical penetration (100 *μ*m or more). Many routine workflows, however, operate on far thinner specimens, such as cultured cell monolayers and tissue sections. Sections used in histopathology and multiplexed antibody-based imaging are typically 5–10 *μ*m thick [2] and seldom more than a few tens of micrometres. For such samples, the added complexity, cost and alignment requirements of these designs still apply, while their characteristic advantages in photodamage and volumetric throughput, which scale with imaging depth, become far less pronounced.

Another approach is to expand the confocal principle to different illumination and detection geometries. Line-scan confocal microscopy focuses the excitation light into a line rather than a diffraction-limited spot [11]. This offers a much higher imaging speed because it completely eliminates one scan direction. The use of a slit aperture introduces a modest asymmetry in the point spread function and slightly reduced axial sectioning compared to point-scanning confocal microscopy, although these effects are generally small under practical operating conditions [12].

Several line-scan confocal implementations have been reported. Early systems employed dedicated line detectors [12], whereas later designs used optical rescanning to project the fluorescence signal onto much more sensitive two-dimensional cameras [13]. Commercial designs are also available. These systems achieve strong performance but have relied on the use of controlled galvanometer mirrors (galvos), which offer limited scanning speeds that usually result in just a single light sweep per image acquisition. They also require more complex synchronization of the mirrors. These limitations not only drive up the complexity of the system but also limit the ability of the system to observe fast events, such as the ‘blinking’ of individual emitters in methods such as single-molecule localization microscopy (SMLM).

We reasoned that a simplified and higher performance detection could be achieved by introducing much faster scanning of the light, using an all-optical synchronization strategy that enables excitation scanning, descanning, and rescanning using a single high-speed scanner. Interestingly, a related concept was already suggested in the original description of line-scan confocal microscopy [11], although in a layout that is difficult to implement in practice.

In this work, we introduce the TriScan, a light-scanning confocal microscope that permits high speed image acquisitions by combining optically-synchronized line confocal imaging with a fast resonant scanner. The TriScan offers a simple and inexpensive optical layout that can be implemented as a compact add-on module for existing microscopes. To the end user, the TriScan presents itself as a widefield microscope with z-sectioning, requiring no custom software or other adjustments to user workflow. We apply the microscope to a variety of samples and show that TriScan enables experiments ranging from high-throughput imaging of large 2D or 3D samples to superresolution measurements using single-molecule localization microscopy (SMLM).

## 2 Results and discussion

Fig. 1A shows an overview of the different confocal modalities developed to date, including the line scanning adopted in our TriScan. A conceptual illustration of this design is shown in Fig. 1B. In this microscope, the excitation light is first shaped into a line using a cylindrical lens. This line is then scanned over the sample using a movable mirror (‘scanner’) positioned in a conjugate back focal plane of the objective. The resulting emission is then descanned (made stationary) by passing over the same movable mirror. A dichroic mirror then separates the emission from the excitation light, passing it through the confocal slit, after which it is reflected by the scanner a third time, sweeping it over a standard imaging camera to build up an image of the sample. We called the device the ‘TriScan’ because the imaging involves three passes of the excitation and emission light over the scanning mirror.

**Figure 1.**
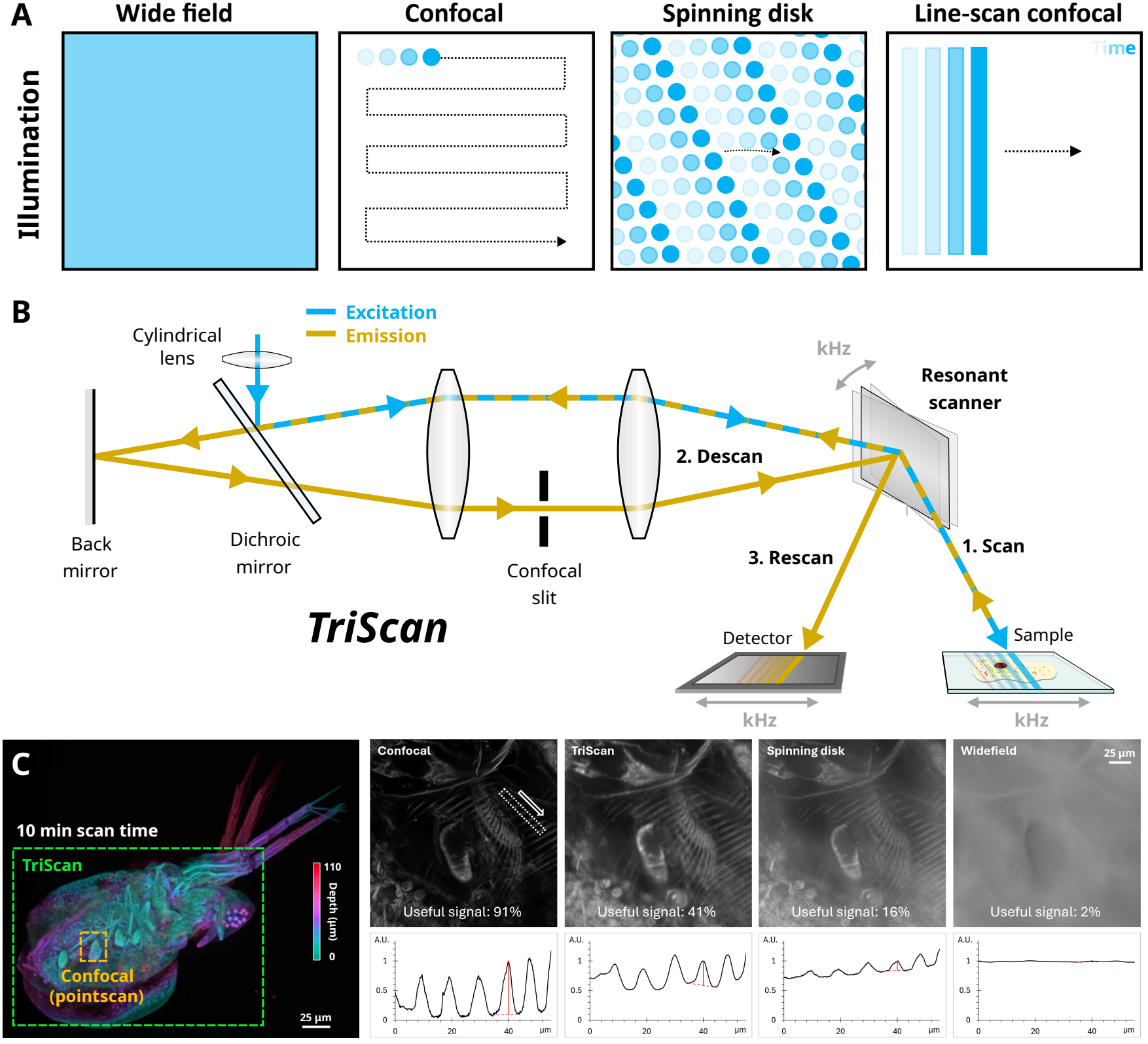
The TriScan concept and practical implementation. **A**, An illustrative comparison of widefield, confocal, spinning disk confocal and line-scan confocal illumination styles. **B**, The optically offset TriScan line-scan confocal implementation: the excitation and emission light are scanned on the sample, descanned through a confocal slit and rescanned on an area detector, using a single resonant scanner while sharing most optical elements for all paths. **C**, TriScan benchmark, using a Daphnia as a sample. Left, a comparison of the area of TriScan or Confocal imaging in 10 minutes, indicating a >50x speedup. Right, a comparison of contrast as quantified in useful signal, for selected techniques.

We explored several designs based on different optical design philosophies, including an optically offset design and an optically centred design, both built as modules suitable for integration into new or existing microscopes. In addition, we have developed a strongly miniaturised, self-contained version with 3D-printed mounts (supplementary information in final publication).

The key defining element of these designs is that they provide all-optical synchronization of the scanning process. In turn, this enables the use of arbitrarily fast scanning mirrors, even if these do not offer full control over the scanner positioning. This includes resonant scanners that readily offer scanning rates up to 16 kHz. In principle, such scanners would allow the TriScan to deliver full field-of-view imaging at a frequency of 32 kHz, though this is very fast compared to the imaging rates of current camera technology and the limited brightness of typical samples. Usually, much slower exposure durations of around tens of milliseconds or more are used instead. With such exposure times, the camera integrates the emission from many light sweeps into a single image. This sweep accumulation is an elegant feature of the TriScan concept, because it increases the chances that fast or transient processes occurring within a single exposure can be reliably detected and sampled.

The sinusoidal movement associated with resonant scanners does result in an inhomogeneous sample illumination because the scanner moves more slowly at the edges of its travel range. Furthermore, for short exposure times streaking artifacts can arise due to the random position of the line at the start of the camera exposure. To compensate for these effects, we introduced a microcontroller that controls the intensity of the excitation light in phase with the motion of the scanner and that takes into account the camera exposure time. This controller is entirely self-contained in that it operates independently from the other hardware and software controlling the measurement, and is therefore invisible to the end user. From an outside perspective, the TriScan offers the ease-of-use and approximates the performance of widefield imaging, while delivering the 3D imaging performance of confocal microscopy.

We next performed a quality assessment of the TriScan and compared it with alternative imaging strategies. We initially imaged a Daphnia sample to compare its performance to that of confocal, spinning disk, and widefield microscopes. The instrumental parameters were optimized to achieve the best performance by trained operators, though we enforced identical exposure times for the TriScan, spinning disk, and widefield modalities. As shown in Fig. 1C, the TriScan outperformed point-scanning confocal in imaging speed by a factor of over 50x, though at the cost of lower contrast. Nevertheless, these images were considerably better than those acquired using widefield and spinning-disk imaging (>20x and >2.5x, respectively). We also investigated the optical quality of the TriScan imaging, revealing that the lateral *x*-resolution of the instrument is essentially identical to that of a classical confocal instrument using the same objective, the *y*-resolution is similar to that of a widefield system since there is no confinement along that axis, and the *z*-resolution is roughly 20 to 30 percent lower than that of a confocal instrument based on a conventional (point-like) aperture. The performance numbers are in line with theoretical estimates [12].

The performance of the TriScan allowed us to image a broad variety of samples, including optically large specimens that would otherwise be time-consuming to image (Fig. 2). One example is a diffuse midline glioma-on-chip model, in which HSJD-DIPG-007 tumour cells and astrocytes are embedded in a ∼750 µm thick hydrogel disk that is supplied with medium only from its edge, so that cellular oxygen consumption establishes a radial gradient from a normoxic periphery to a hypoxic core [14] (Fig. 2A). Imaging across this millimetre-scale sample allows cell distribution and behaviour to be compared along the entire oxygen gradient. We further imaged several volumetric multicolour mouse samples (Fig. 2B-D). A mammary gland from a genetic mouse model was used to compare the clonal growth of PIK3CA(H1047R) mutant cells with that of recombined wild-type neighbours, following sporadic Cre-mediated recombination induced by intraductal Tat-Cre injection (Fig. 2C). A second, more resolution-demanding sample was a 3D culture of primary rat neurons grown in networks with astrocytes. Dense synaptic contacts between neurons were revealed using a synapsin antibody (Fig. 2D). Finally, we imaged a ∼ 140 µm thick mouse colon sample labelled for nuclei (DAPI), vimentin, blood vessels (*Lycopersicon esculentum* lectin, LEL) and enteric neurons (PGP9.5) (Fig. 2B), allowing the enteric nervous system to be visualised in the context of the surrounding tissue architecture. Across all of these applications, the TriScan allowed us to quickly acquire high resolution images.

**Figure 2.**
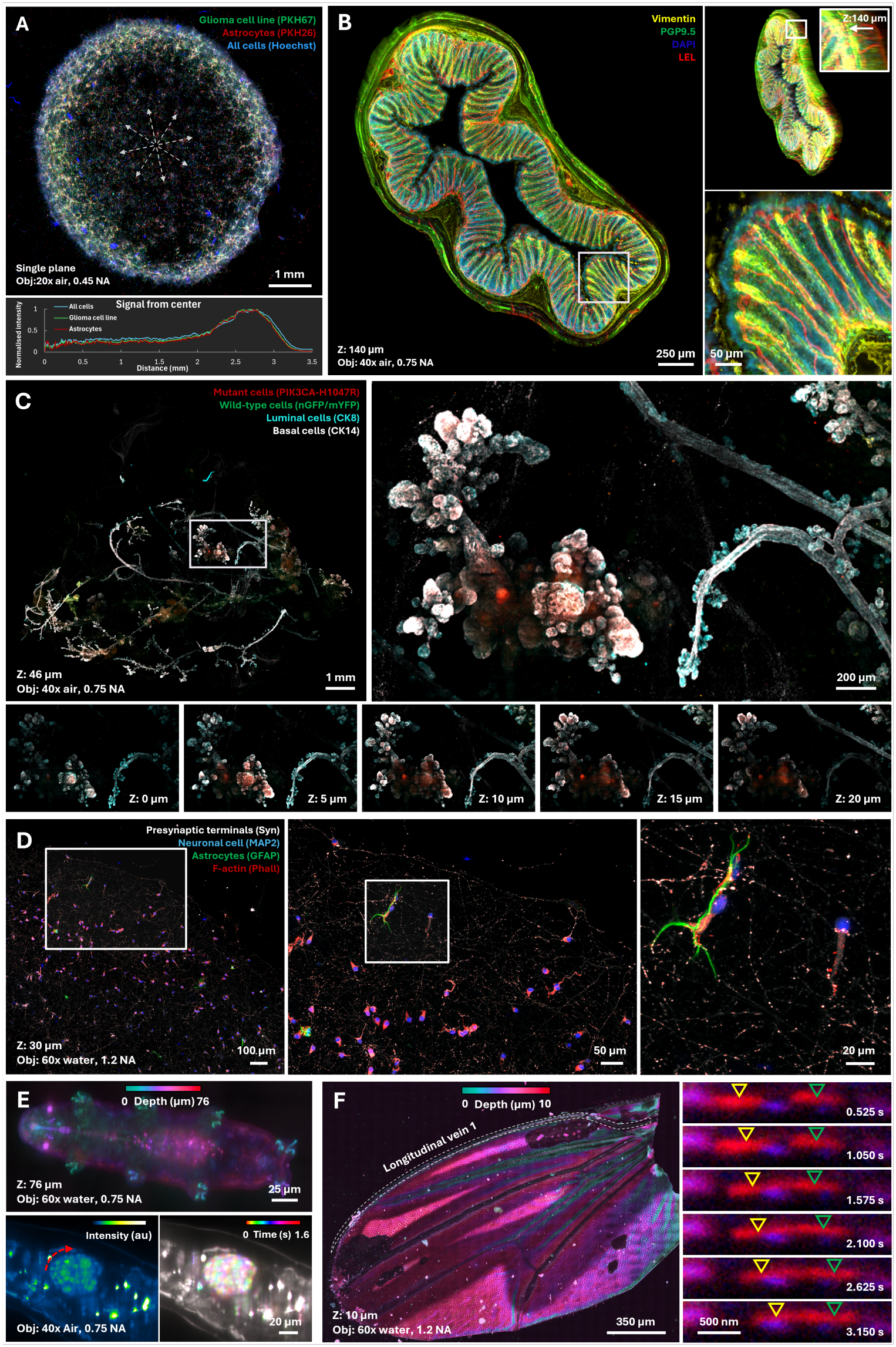
Volumetric imaging of diverse biological samples with the TriScan. **A**, DMG-on-Chip model of PKH67-labelled HSJD-DIPG-007 cells (green), PKH26-labelled human brainstem astrocytes (red) and all cells (Hoechst, blue): a single optical section through the hydrogel disk, which is subject to a radial oxygen gradient from a normoxic periphery to a hypoxic core; the plot shows radial intensity profiles from the disk centre. **B**, 140 µm thick mouse colon sample labelled for nuclei (DAPI), vimentin, blood vessels (LEL) and enteric neurons (PGP9.5). **C**, Mammary gland of a Red2Onco-PIK3CA(H1047R) mouse, showing PIK3CA(H1047R) mutant cells (RFP), recombined wild-type epithelial cells (nGFP/mYFP), luminal cells (CK8) and basal cells (CK14). **D**, 3D culture of primary rat neurons labelled for neuronal cells (MAP2), astrocytes (GFAP), F-actin (phalloidin) and presynaptic terminals (synapsin). **E**, Live tardigrades: full-body image of an ethanol-immobilised specimen (autofluorescence) and time-lapse imaging of a freely moving animal. **F**, Mitochondria moving along the longitudinal vein of the wing of a transgenic *Drosophila* expressing mitochondria-targeted GFP.

The speed of TriScan imaging also lends itself well to the visualization of dynamic samples. We explored this via the whole-organism imaging of sedated tardigrades and by revealing fast intra-organismal dynamics (Fig. 2E). We also imaged anaesthetized *Drosophila* mounted on the instrument and tracked the 3D movement of mitochondria along the longitudinal vein (L1) of the wing. In both cases, well-sampled 3D dynamic imaging was made possible by the combination of speed and high-contrast imaging inherent to TriScan (Fig. 2F). An earlier prototype likewise was successfully applied to linear dichroism-based biosensing within live organisms [15].

We next explored 3D super-resolution imaging using two distinct methods. In super-resolution optical fluctuation imaging (SOFI), the sample is labelled with blinking fluorophores that are then visualized by rapidly acquiring a stack of multiple images. The resulting image stack is then subjected to a statistical analysis to produce a super-resolution image [16, 17]. The advantage of this method is that it can operate under a broad range of conditions, including low emitter brightness or high emitter densities [18, 19]. We found that the TriScan could be used to acquire high-resolution SOFI images on living cells labelled with a HaloTag (Fig 3A).

**Figure 3.**
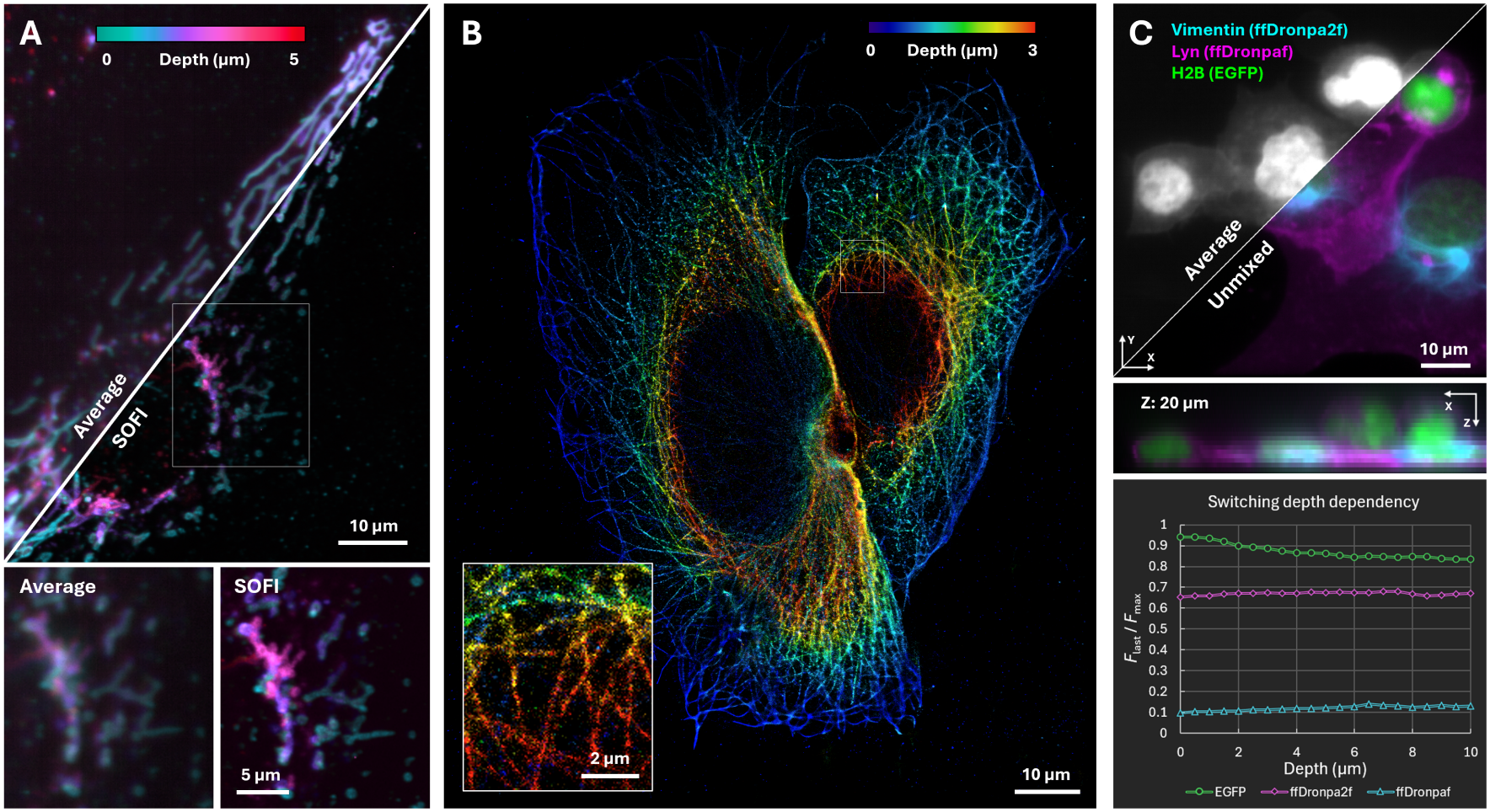
Super-resolution and multiplexed imaging with the TriScan. **A**, SOFI of live COS-7 cells expressing DAKAP-HaloTag (mitochondrial outer membrane) labelled with JF635b, acquired at 5 axial planes and analysed with second-order cross-cumulants. **B**, DNA-PAINT image of microtubules (*α*-tubulin) in fixed U2OS cells, acquired at 7 axial planes and rendered from the localised single-molecule events. **C**, Switching-kinetics unmixing of three green-emitting labels in fixed COS-7 cells: H2B-EGFP (chromatin), Lyn-ffDronpaF (plasma membrane) and vimentin-ffDronpa2F (vimentin filaments), separated by their response to a repeating 405/488 nm illumination sequence.

We also applied the microscope to SMLM-based imaging using DNA-PAINT. Traditionally, this methodology makes use of widefield imaging to image many fluorophores in parallel and to detect emission events that may appear or disappear at any time during an exposure. However, the lack of optical sectioning in this approach has complicated the application of this methodology to larger samples, often limiting it to total internal reflection (TIRF) or highly inclined illumination [20]. Several approaches have been introduced to extend SMLM to larger samples, based on, for example, spinning-disk, confocal, or light-sheet illumination [21–24], often requiring complex instrumentation. The TriScan, in contrast, combines optical sectioning with a simple, camera-based architecture in which many light sweeps are integrated per camera exposure, increasing the probability that transient binding events are detected. Overall, this enabled the DNA-PAINT imaging of thicker samples (Fig 3B).

Finally, we used the TriScan for multiplexed imaging based on the optical switching of reversibly switchable fluorescent proteins. These fluorophores can be switched between a fluorescent and a non-fluorescent state by light, differing in the efficiency with which they respond [25–27]. Labels with strongly overlapping spectra can therefore be distinguished by their response to a defined illumination sequence rather than by their colour. This methodology is typically performed in widefield imaging due to the need to accumulate multiple observations at each position on the sample, preventing the routine application of confocal-based methods. The combination of temporally-modulated excitation and the fast imaging performance of the TriScan allowed us to readily separate three green-emitting labels, H2B-EGFP, Lyn-ffDronpaF and vimentin-ffDronpa2F, targeted to chromatin, the plasma membrane and vimentin filaments in fixed COS-7 cells (Fig 3C).

## 3 Conclusion

In this work, we introduced the TriScan, a fluorescence microscope that uses all-optical synchronization in combination with very fast scanning mirrors to deliver confocal imaging with the ease of use and performance of widefield imaging. Our design enables high imaging speeds, sensitive fluorescence detection, and straightforward operation within a compact and inexpensive optical layout. Owing to its conceptually simple design and modest hardware requirements, the materials cost specific to TriScan is below 10,000 euros, which is small compared to the costs associated with the components intrinsically required for fluorescence imaging, including the excitation light source, microscope body, objective, and camera.

We demonstrated the versatility of this design across a range of imaging modalities, including multicolour, 2D, 3D, dynamic, and superresolution imaging. We implemented the system as an add-on module for existing fluorescence microscopes, allowing confocal imaging capabilities to be added without modifications to acquisition software or user workflows. From the operator’s perspective, the system behaves as a conventional widefield microscope. The simplicity of the optical concept also permits its adaptation to a wide range of existing and new instrumentation.

By combining performance, simplicity, and affordability, the TriScan lowers the barrier to fast 3D fluorescence imaging and can provide a practical alternative to more complex confocal imaging platforms. We anticipate that the approach will facilitate the development of robust and cost-effective instruments for applications ranging from fundamental research to diagnostic imaging and high-content screening. More broadly, the combination of high-speed acquisition, optical sectioning, implementation flexibility, and compatibility with existing microscope platforms establishes this method as a versatile and accessible approach for fluorescence microscopy across a wide range of imaging applications.

## 4 Methods

### TriScan specifics

All imaging was performed on the optically offset TriScan, unless stated otherwise. Both TriScan configurations used a quad-band dichroic and emission filter set for the 405, 488, 561 and 640 nm laser lines (ZET405/488/561/635, 25 mm, Chroma), an Oxxius laser combiner (405, 488, 561 and 640 nm) and an SC-30 resonant scanner (EOPC). The optically offset TriScan was equipped with an ORCA-Fusion BT digital CMOS camera (Hamamatsu) and the optically centred TriScan with a ZL41 Cell sCMOS camera (Andor, Oxford Instruments). The following Olympus objectives were used: 60× water immersion, 1.2 NA (UPLSAPO60XW); 40× air, 0.75 NA (UPLFLN); 20× air, 0.45 NA (LUCPLFLN). Unless stated otherwise, the confocal slit width corresponded to 1.15 Airy units on the optically offset TriScan and 0.89 Airy units on the optically centred TriScan.

### Tardigrades

Tardigrades (species indet.) were collected on 26 May 2025 (10:40) from moss growing on tree bark in a wooded area along Kapeldreef, Heverlee (Leuven), Belgium (50.8624° N, 4.6713° E, WGS84; elevation ∼27 m a.s.l.). Animals were extracted by soaking the moss in tap water overnight, transferring the water to a Petri dish and screening it for tardigrades under a standard student-grade compound brightfield microscope (Kentfaith GW45.0066). All specimens were imaged (Fig. 2E) within 36 h of collection. The specimen shown in full was sedated in 7% (v/v) ethanol in tap water during imaging. The full-body image was acquired with 488 nm excitation, a 60× water-immersion objective (1.2 NA) and 300 ms exposure time; the dynamics were imaged with 561 nm excitation, a 40× air objective (0.75 NA) and 100 ms exposure time.

### Mouse mammary gland

Mammary gland tissue (Fig. 2C) was obtained from an adult female Red2Onco-PIK3CA(H1047R) mouse [28]. In this model, sporadic Cre-mediated recombination stochastically activates one of four fluorescent reporters per cell: tdimer2(12) RFP co-expressed with oncogenic PIK3CA(H1047R) (mutant cells), or, in wild-type cells, nuclear GFP (nGFP), membrane-tethered mCerulean (mCFP) or monomeric EYFP (mYFP A206K, cytosolic and nuclear). Mutant cells and recombined wild-type neighbours can therefore be visualised in the same tissue, while non-recombined cells remain unlabelled. Recombination was achieved by mammary intraductal injection of TAT-Cre recombinant protein (20 U per gland), and glands were harvested 28 days after recombination. All animal procedures were approved by the Animal Welfare Committee of KU Leuven (ethical approval numbers P155/2020 and Onco-088/2025).

Dissected mammary glands were incubated in HBSS (Gibco) containing collagenase A (2.5 mg/ml; Sigma-Aldrich, 10103578001) and hyaluronidase (50 µg/ml; Sigma-Aldrich, H3506) for 25 min at 37 °C with shaking (180 rpm). Tissues were subsequently fixed in periodate-lysine-paraformaldehyde (PLP) buffer (1% PFA (Electron Microscopy Sciences), 0.01 M sodium periodate, 0.075 M L-lysine and 0.0375 M phosphate buffer (0.081 M Na_2_HPO_4_, 0.019 M NaH_2_PO_4_; pH 7.4)) for 2 h at room temperature on an orbital shaker (25 rpm). Glands were then incubated for at least 3 h in blocking buffer containing 1% bovine serum albumin (Roche Diagnostics), 5% normal goat serum (Monosan) and 0.8% Triton X-100 (Sigma-Aldrich) in PBS. Primary antibodies were diluted in blocking buffer and incubated overnight at room temperature; secondary antibodies, diluted in blocking buffer, were incubated for at least 6 h. Luminal epithelial cells were immunostained for cytokeratin 8 (CK8; rat, clone TROMA-1, Sigma-Aldrich, MABT329; 1:500) with Alexa Fluor 405 goat anti-rat (Thermo Fisher Scientific, A48261; 1:400), and basal cells for cytokeratin 14 (CK14; rabbit, Covance, PRB-155P; 1:600) with Alexa Fluor 647 goat anti-rabbit (Thermo Fisher Scientific, A21245; 1:400). Glands were washed with PBS and mounted on a microscope slide in Vectashield HardSet (Vector Laboratories, H-1400).

Mutant cells (RFP) were excited at 561 nm, recombined wild-type cells (nGFP/mYFP) at 488 nm, CK8 at 405 nm and CK14 at 640 nm, using a 40× air objective (0.75 NA) and 35 ms exposure time. nGFP and mYFP were not spectrally separated, and mCFP also contributes to the 405 nm channel. To correct for bleed-through of the Alexa Fluor 647 signal into the 561 nm channel, the 640 nm image was scaled by a factor of 1/2.5 and subtracted from the 561 nm image. The scaling factor was estimated from regions that should only contain Alexa Fluor 647 signal, by comparing their intensity in the 640 nm and 561 nm channels.

### Neurons

Primary hippocampal neurons were isolated from E18 rat embryos and cultured in 3D in a Matrigel/collagen hydrogel for 21 days in vitro (Fig. 2D). Cultures were fixed with 4% PFA for 20 min, permeabilised with 0.1% Triton X-100 and blocked with 0.25% fish gelatin. Neurons were labelled with chicken anti-microtubule-associated protein 2 (MAP2; Abcam, #ab5392, 1:5000) followed by a custom CF405-conjugated anti-chicken secondary antibody (Biotium), astrocytes with an ATTO 488-labelled single-domain antibody against glial fibrillary acidic protein (GFAP; Synaptic Systems), actin with Alexa Fluor 568 phalloidin (Thermo Fisher Scientific, #A12380), and presynaptic terminals with rabbit anti-synapsin 1 (Synaptic Systems, #106 008) followed by an Alexa Fluor 647-conjugated donkey anti-rabbit IgG (H+L) highly cross-adsorbed secondary antibody (Thermo Fisher Scientific). Samples were mounted on glass slides in ProLong Gold mounting medium.

Imaging was performed using a 60× water-immersion objective (NA 1.2) with an exposure time of 20 ms. The confocal slit width corresponded to 0.86 Airy units. To correct for bleed-through from the 640 nm into the 561 nm channel, the 640 nm image was scaled by a factor of 1/2.5 and subtracted from the 561 nm image. The scaling factor was estimated from regions that should only contribute to the 640 channel, by comparing their intensity in the 640 nm and 561 nm channels.

### Oxygen gradient chamber

HSJD-DIPG-007 cells and human brainstem astrocytes (ScienCell) were cultured in a DMG-on-Chip (DoC) device as described previously [14]. Prior to seeding, HSJD-DIPG-007 cells and astrocytes were labelled with PKH67 (green; Sigma-Aldrich, PKH67GL) and PKH26 (red; Sigma-Aldrich, PKH26GL), respectively, to discriminate the two cell types. For each cell population, 1 µl of PKH stock solution was diluted in 250 µl Diluent C, and 50 µl of this solution was mixed with a further 50 µl Diluent C. Cells were washed with PBS, pelleted, resuspended in the labelling solution and incubated for 5 min at room temperature. Labelling was stopped by adding tumour stem medium (TSM) supplemented with 1% BSA, after which cells were centrifuged (400× *g*, 5 min) and resuspended in fresh TSM.

Both cell types were seeded at 5 000 cells/µl each (1:1 ratio; 10 000 cells/µl in total) in a hydrogel consisting of 90% Matrigel (Corning, 352637) and 10% high-molecular-weight hyaluronic acid (5 mg/ml; R&D Systems). The cell-containing matrix was deposited in the centre of a pre-shaped PDMS spacer on a 40× 60 mm glass coverslip (Epredia, #1, 0.13–0.16 mm), and a second coverslip was placed on top to flatten the matrix into a uniform disk, surrounded by a medium-filled gutter. Because medium and oxygen are supplied only from the edge of the disk and cellular oxygen consumption exceeds diffusion, a radial oxygen gradient is established from near-normoxic conditions at the periphery to hypoxia in the core [14].

After 48 h of culture in TSM at 37 °C and 5% CO_2_, the upper coverslip was removed and nuclei were stained with Hoechst 33342 (Thermo Fisher Scientific, 33342; 1 µg/ml) for 1 h. Samples were rinsed three times with PBS, fixed in freshly prepared 2% paraformaldehyde and 0.1% glutaraldehyde in PBS for 20 min at room temperature, washed three times with PBS, mounted in Fluoromount-G (Invitrogen, 00-4958-02) and covered with a glass coverslip.

This sample (Fig. 2A) was imaged on the optically centred TriScan using a ×20 air objective (0.45 NA), with Hoechst excited at 405 nm, PKH67 at 488 nm and PKH26 at 561 nm, and 50 ms exposure time.

Radial intensity profiles were calculated for each channel separately using a custom analysis, which averages pixel intensities in concentric rings of 5 pixels width around the approximate centre of the sample, using a circular selection that enclosed the entire sample and extended slightly beyond its edge.

### SOFI

COS-7 cells (ATCC CRL-1651) were cultured in DMEM supplemented with 10% FBS and GlutaMAX at 37 °C and 5% CO_2_. Cells (7 ×10^4^) were seeded in a 35 mm glass-bottom dish (MatTEK p35g-1.5-14-c). After 24 h, cells were transfected with 2 µg pcDNA3-DAKAP-HaloTag using FuGENE 6 (Promega) at a 1:3 DNA:reagent ratio, following the manufacturer’s instructions. After a further 24 h, cells were labelled overnight with 500 nM JF635b-HaloTag ligand, washed once with PBS and imaged live in HBSS at RT.

Imaging (Fig 3A) was performed with 640 nm excitation and a 60× water-immersion objective (1.2 NA). At each of 5 axial planes, 500 frames were acquired with 60 ms exposure time. SOFI analysis was performed in Localizer [29] using second-order cross-cumulants.

### DNA-PAINT

U2OS cells (ATCC HTB-96) were cultured in DMEM supplemented with 10% FBS at 37 °C and 5% CO_2_. Cells (1.5 ×10^4^) were seeded in 8-well glass-bottom µ-Slides (#1.5H, Ibidi). After 24 h, cells were fixed with prewarmed 4% PFA in PBS (pH 7.4) for 30 min, and unreacted aldehydes were quenched with 0.1 M glycine in PBS for 15 min. Cells were permeabilised and blocked for 30 min at RT in 3% BSA/0.25% Triton X-100 in PBS, incubated with mouse anti-*α*-tubulin (Synaptic Systems, 302 211; 2 µg/ml in blocking buffer) for 1 h at RT, washed 3 × 5 min with PBS, and incubated with MASSIVE-AB 3-PLEX anti-mouse secondary nanobody carrying docking strand 1 (Massive Photonics; 5 µg/ml) for 1 h at RT. After washing 3 × 5 min, cells were imaged in Imaging Buffer (Massive Photonics) containing 5 nM Imager 1-Cy3B and gold nanoparticles (Massive Photonics) as fiducial markers.

The single-molecule data for Fig 3B comprised 20 000 frames per plane for 7 axial planes, acquired with 561 nm excitation, 100 ms exposure time and a confocal slit width corresponding to 0.86 Airy units. Raw data was localised using custom software [30] and drift-corrected within each plane using COMET [31]. In order to assemble the 7 planes into a volumetric image, the localisations (camera pixel size 104.45 nm) were processed with a custom Igor Pro procedure which registered consecutive planes by cross-correlation of their localisation density images. For display purposes, localisations from fiducial beads were excluded using a density-based criterion, with remaining bead signal masked manually. Each localisation was rendered as a 2D Gaussian (σ = 8 nm) on a 5 nm pixel grid and exported as 16-bit TIF images.

### Switching-kinetics unmixing

COS-7 cells (ATCC CRL-1651) were cultured in DMEM supplemented with 10% FBS at 37 °C and 5% CO_2_. Cells (7 × 10^4^) were seeded in 35 mm glass-bottom dishes (MatTEK p35g-1.5-14-c) and transfected with H2B-EGFP, pcDNA3-Lyn-ffDronpaF and pcDNA3-vimentin-ffDronpa2F (666 ng per plasmid) using FuGENE 6 (Promega) (at a 1:3 DNA:reagent ratio), following the manufacturer’s instructions. Single-plasmid transfections, used to obtain reference switching kinetics, and a triple co-transfection were prepared in parallel. After 24 h, cells were fixed with 4% PFA in PBS (pH 7.4) for 30 min, quenched with 0.1 M glycine in PBS for 15 min, washed with PBS and imaged in PBS.

Imaging (Fig. 3C) was performed with a repeating illumination sequence of one dark frame (no laser), two 405 nm frames, a second dark frame and ten 488 nm frames, each with an exposure time of approximately 10 ms, and a confocal slit width corresponding to 0.86 Airy units. The three labels were unmixed per pixel using an adaptation of Chain and Conquer [32]. Custom software was used for the analysis.

### Mouse colon and *Daphnia*

Both samples were obtained commercially: the mouse colon sample from Luxidea (OptiSlides) and the whole-mount *Daphnia* from AmScope (PS100B General Biology glass slide set). The mouse colon sample (∼140 µm thick; Fig. 2B) was labelled for DAPI, vimentin, PGP9.5 and LEL and imaged with a 40× air objective (0.75 NA), 405, 488, 561 and 640 nm excitation and 25 ms exposure time. The *Daphnia* (Fig. 1C) was imaged with a 60 ×water-immersion objective (1.2 NA), 25 ms exposure time and 488 nm excitation.

For the comparison with commercial instruments (Fig. 1C), we used an Olympus IX71 widefield microscope with Lumencor illumination (Arenberg Optical Imaging Center, KU Leuven), a Nikon Ti2 with a CrestOptics CICERO spinning-disk unit (pinhole: 1.49 Airy units; BioImaging Core Leuven, VIB, KU Leuven) and a Leica TCS SP8 X confocal microscope (pinhole: 1 Airy unit; Hofkens lab, KU Leuven). Useful signal is quantified as the peak-to-valley ratio of the highlighted peaks in the line profiles, with the peak fluorescence normalised to 1.

### *Drosophila* wing

Transgenic *Drosophila melanogaster* expressing mitochondria-targeted GFP (*UAS-mitoGFP; D42-GAL4*) were raised on standard food at room temperature under a natural light/dark cycle. Adult flies were immobilised using a dedicated mounting setup, in which the animal was positioned between a coverslip and an agar gel pad (H. Roberge et al., in preparation). The wings were gently isolated using a fine brush and placed in a small droplet of glycerol to facilitate imaging, and flies were anaesthetised with CO_2_ on the microscope to minimise movement. Mitochondrial dynamics were imaged in vivo within the longitudinal wing veins of living animals. Wing overview images (Fig. 2F) were acquired after live imaging, on the desiccated wing.

Imaging specifics: All *Drosophila* imaging was performed with 488 nm excitation and a 60× water-immersion objective (1.2 NA), at 25 ms exposure time over a range of 21 depth planes (500 nm stepsize).

## Acknowledgements

The authors gratefully acknowledge Luke Lavis for providing the JF635b-HaloTag ligand, Bon-Kyoung Koo and Benjamin D. Simons for providing the Red2Onco-PIK3CA(H1047R) mouse line, and Angel Montero Carcaboso for providing the HSJD-DIPG-007 cell line. The authors also thank Michel Sliwa for his continued support and valuable discussions, Jean-Emmanuel Clement for his assistance with imaging in the oxygen gradient chamber, and Lea Mureta for her help with tardigrade collection.

## Funding

This work received funding from the Research Foundation Flanders (FWO) under grants 1S55225N, 11L7222N, 12B2Y24N, 12A7225N, the KU Leuven under grant PDMT2/23/043, and the Deutsche Forschungsgemeinschaft under grant 560074072. Additionally, this work received funding from the FWO under grants G010723N and I009022N, the KU Leuven under grant STG/22/059, the French National Research Agency (ANR) under grant ANR-19-CE16-0012, FLAG-ERA under grant SENSEÏ ANR-19-HBPR-0003, the European Union’s European Innovation Council (EIC) Pathfinder Open Programme under grants CHEM-SCAN 101259077, UniSens 101131014, the European Research Council (ERC) under grant 101113439, the Initiative d’Excellence of Université de Lille under grant WILL-CHAIRES-22-001-DEEPSEE, de Stichting tegen Kanker under grant F/2024/2642, and the Kom op Tegen Kanker Emmanuel van der Schueren Finisher Grant.

## Conflict of interest

RVDE, WV, and PD have a patent on one of the instrumental designs (EP4437377, WO2023/094519A1). JL is independently involved in commercializing instruments based on this concept via Innovative Bioimaging.

## Notes

### Summary of Updates

Added additional measurements and characterization; Authors updated; Figures revised

## References

1. Vandereyken, K., Sifrim, A., Thienpont, B. & Voet, T. Methods and applications for single-cell and spatial multi-omics. Nat. Rev. Genet. 24, 494–515 (2023).

2. Hickey, J. W. et al. Spatial mapping of protein composition and tissue organization: a primer for multiplexed antibody-based imaging. Nat. Methods 19, 284–295 (2022).

3. Bray, M.-A. et al. Cell Painting, a high-content image-based assay for morphological profiling using multiplexed fluorescent dyes. Nat. Protoc. 11, 1757–1774 (2016).

4. Leung, C. M. et al. A guide to the organ-on-a-chip. Nat. Rev. Methods Primers2, 33 (2022).

5. Handbook of Biological Confocal Microscopy 3rd ed. (ed Pawley, J. B.) (Springer, New York, 2006).

6. Galland, R. et al. 3D high- and super-resolution imaging using single-objective SPIM. Nat. Methods 12, 641–644 (2015).

7. Dunsby, C. Optically sectioned imaging by oblique plane microscopy. Opt. Express 16, 20306–20316 (2008).

8. Bouchard, M. B. et al. Swept confocally-aligned planar excitation (SCAPE) microscopy for high-speed volumetric imaging of behaving organisms. Nat. Photonics 9, 113–119 (2015).

9. Sapoznik, E., Chang, B.-J., et al. A versatile oblique plane microscope for largescale and high-resolution imaging of subcellular dynamics. eLife 9, e57681 (2020).

10. Yang, B., Lange, M., Millett-Sikking, A., et al. DaXi—high-resolution, large imaging volume and multi-view single-objective light-sheet microscopy. Nat. Methods 19, 461–469 (2022).

11. Brakenhoff, G. J. & Visscher, K. Confocal imaging with bilateral scanning and array detectors. J. Microsc. 165, 139–146 (1992).

12. Wolleschensky, R., Zimmermann, B. & Kempe, M. High-speed confocal fluorescence imaging with a novel line scanning microscope. J. Biomed. Opt. 11, 064011 (2006).

13. Lee, J., Miyanaga, Y., Ueda, M. & Hohng, S. Video-rate confocal microscopy for single-molecule imaging in live cells and superresolution fluorescence imaging. Biophys. J. 103, 1691–1697 (2012).

14. Terrassoux, L. et al. Novel Diffuse Midline Glioma-on-Chip Recapitulating Tumor Biophysical Microenvironment to Assess the Heterogeneity of Response to Therapies. Small 21, e05343 (2025).

15. Miclea, P. et al. FLIPs: Genetically encoded molecular biosensors for functional imaging of cell signaling by linear dichroism microscopy. Sci. Adv. 12, eadz5662 (2026).

16. Dertinger, T., Colyer, R., Iyer, G., Weiss, S. & Enderlein J. Fast, background-free, 3D super-resolution optical fluctuation imaging (SOFI). Proc. Natl. Acad. Sci. U. S. A. 106, 22287–22292 (2009).

17. Dedecker, P., Mo, G. C. H., Dertinger, T. & Zhang, J. Widely accessible method for superresolution fluorescence imaging of living systems. Proc. Natl. Acad. Sci. U. S. A. 109, 10909–10914 (2012).

18. Geissbuehler, S., Dellagiacoma, C. & Lasser, T. Comparison between SOFI and STORM. Biomed. Opt. Express 2, 408–420 (2011).

19. Vandenberg, W., Leutenegger, M., Duwé, S. & Dedecker, P. An extended quantitative model for super-resolution optical fluctuation imaging (SOFI). Opt. Express 27, 25749–25766 (2019).

20. Tokunaga, M., Imamoto, N. & Sakata-Sogawa, K. Highly inclined thin illumination enables clear single-molecule imaging in cells. Nat. Methods 5, 159–161 (2008).

21. Schueder, F. et al. Multiplexed 3D super-resolution imaging of whole cells using spinning disk confocal microscopy and DNA-PAINT. Nat. Commun. 8 (2017).

22. Thiele, J. C. et al. Confocal Fluorescence-Lifetime Single-Molecule Localization Microscopy. ACS Nano 14, 14190–14200 (2020).

23. Oleksiievets, N. et al. Fluorescence lifetime DNA-PAINT for multiplexed super-resolution imaging of cells. Communications Biology 5 (2022).

24. Cheng, S., Nakatani, Y., Gagliano, G., Saliba, N. & Gustavsson, A.-K. Light sheet illumination in single-molecule localization microscopy for imaging of cellular architectures and molecular dynamics. npj Imaging 2 (2024).

25. Valenta, H. et al. Separation of spectrally overlapping fluorophores using intraexposure excitation modulation. Biophys. Rep. 1, 100026 (2021).

26. Valenta, H., Dalla Vecchia, M., Van den Eynde, R., Vandenberg, W. & Dedecker, P. Super-resolved Imaging of Molecular Interactions Using FRET-SOFI. Nano Lett. 25, 16690–16696 (2025).

27. Duwé, S., Vandenberg, W. & Dedecker, P. Live-cell monochromatic dual-label sub-diffraction microscopy by mt-pcSOFI. Chem. Commun. 53, 7242–7245 (2017).

28. Yum, M. K. et al. Tracing oncogene-driven remodelling of the intestinal stem cell niche. Nature 594, 442–447 (2021).

29. Dedecker, P., Duwé, S., Neely, R. K. & Zhang, J. Localizer: fast, accurate, open-source, and modular software package for superresolution microscopy. J. Biomed. Opt. 17, 126008 (2012).

30. Bates, M., Jones, S. A. & Zhuang, X. in Imaging: A Laboratory Manual (ed Yuste, R.) 547–576 (Cold Spring Harbor Laboratory Press, Cold Spring Harbor, NY, 2011).

31. Reinkensmeier, L., Aufmkolk, S., Farabella, I., Egner, A. & Bates, M. Cost-function Optimized Maximal Overlap Drift Estimation for Single Molecule Localization Microscopy. bioRxiv. Preprint (2026).

32. Gómez-Sánchez, A. et al. Chain and Conquer: blind kinetic unmixing of reversibly switchable fluorescent proteins. Anal. Chem. Manuscript under review (2026).

